# Inbreeding and demography interact to impact population recovery from bottlenecks

**DOI:** 10.1101/2024.12.10.627850

**Authors:** Jia Zheng, Ella Rees-Baylis, Thijs Janzen, Zhengwang Zhang, Xiangjiang Zhan, Daiping Wang, Xiang-Yi Li Richter

## Abstract

Biodiversity loss driven by climate change and human activities poses a critical global challenge. Population restoration and reintroduction programs are essential for mitigating this threat, yet their outcomes are often unpredictable due to poorly understood success factors. The conservation program of the crested ibis (*Nipponia nippon*) marks a successful example where the population rose from seven survivors to over 9,000 in the past four decades. To learn whether this successful restoration was due to chance or largely repeatable, we developed an individual-based model that simulates the restoration process by incorporating life-history parameters from empirical data. Our simulation results closely mirror empirical findings, including the time taken to reach the current population size and population-level inbreeding coefficients. We further analyzed the model to compare the effectiveness of two reintroduction strategies and analyzed how inbreeding depression interacts with demography to influence the chance of recovery from bottlenecks. The reintroduction simulations reveal that the ‘firework’ approach (one-source translocations) outperforms the ‘stepping-stone’ (serial translocations) approach in restoration effectiveness. Our simulations over broad demographic parameters demonstrate that the net effect of inbreeding varies with species-specific demography, and highlight the importance of considering this interaction when interpreting conservation outcomes and designing future reintroduction programs.

## Introduction

More than one in four species assessed by the IUCN is threatened with extinction, and biodiversity is predicted to further decline as the effects of climate change and anthropomorphic threats increase^1^. To reverse this trend, effective conservation programs are in pressing need. Predicting the likelihood of population recovery from ecological threats and providing optimal species reintroduction strategies are therefore central tasks in these efforts^2, 3, 4, 5^. Despite the evident need, the outcomes of population restoration and reintroduction across different species are highly variable^3, 6, 7, 8, 9, 10^, and the causes of failures are often not fully understood^9, 11, 12^. Subsequent in-depth analyses of such empirical conservation outcomes can help us better understand potential causes underlying (un)successful outcomes and draw broader conclusions that can be applied to conservation efforts for other endangered species^3, 5, 12, 13^.

Bottlenecked populations can result from ecological catastrophes, yet also reintroduction programs often inevitably create small populations due to financial constraints, conservation uncertainties, and to avoid negative impacts on the source population^7, 10, 14^. The primary goals of such programs are to help endangered species maintain genetic diversity by establishing multiple self-sustaining populations or to help recover their geographic distributions to historical scales^5, 14, 15^. However, unavoidable inbreeding in small populations during the initial phase of reintroduction often results in a loss of genetic diversity, which raises looming concerns about long-term conservation success^3, 11^. Inbreeding is therefore likely to play a role in the variable outcomes of restoration efforts.

Inbreeding depression, the reduced fitness of a population due to mating between genetically close-related individuals, is regarded as one of the most cryptic threats to population viability and has recently garnered increasing research attention^2, 16, 17^. In wild animals, inbreeding is known to result in abnormal embryo development^18, 19^ and reduced fecundity^16^, survival rates^2, 20, 21^, and recruitment probability^22, 23^. These effects stem from decreased genetic variation, which can result in the accumulation of rare recessive deleterious alleles. Despite this, inbreeding is common in many animal species, with little evidence of avoidance of mating with kin^24^. Genomic-wide sequencing has identified that considerable inbreeding persists even in species that successfully recovered from bottleneck events^2, 13, 25, 26^. There is thus some concern that the detrimental effects of inbreeding could somehow undermine the viability of current steadily growing populations, even leading to a future collapse during reintroduction programs^13, 16, 25, 27^.

A remarkable conservation success is that of the crested ibis (*Nipponia nippon*) in East Asia. This long-lived species faced a severe bottleneck in the last century due to habitat loss and pollution^28^. Fortunately, with extensive conservation efforts, its population has been officially announced to have rebounded from just seven wild individuals to over 9000 (wild and captive) individuals in the past four decades^29, 30^. Like many reintroduction programs, it remains unclear how and why this species has overcome the detrimental impact of inevitable inbreeding and what lessons can be learned to guide conservation efforts for other endangered species.

The crested ibis is thus an ideal empirical example for which a more extensive analysis of its conservation can be of great use, especially given that long-lived species are thought to be particularly vulnerable to extinction during bottleneck events^13, 16^. For instance, their slow reproductive pace may hinder them to rapidly recover from a small population size^3, 13^. Despite this, negative fitness consequences from bottleneck events appear to be strikingly negligible in some long-lived critically endangered species^6, 7, 13, 16, 17, 31^. This has partially been attributed to the idea that harmful effects of inbreeding can potentially be buffered by repeated reproduction over a long lifespan, which aids in successful restoration^16, 17, 32^. In other words, lifetime fecundity can be substantial for species that have a slow birth rate yet continue to breed over decades. However, the influence of demography such as fecundity and mortality rates are often overlooked when predicting the effects of inbreeding on population viability^2, 5, 13, 21^.

Therefore, in this study, we created an individual-based model that incorporates both empirical inbreeding depression data and demographic rates from the crested ibis population restoration. We aimed first to investigate to what extent such a theoretical model can make predictions that align with empirical conservation outcomes and to understand whether the crested ibis restoration success was achieved by chance or is largely repeatable. We then conducted two additional model analyses, as examples of broader lessons we can learn from the successful conservation efforts. The first analysis uses the same ibis system to compare the efficiency of two idealized reintroduction approaches: a ‘firework’ approach, where founders from the originally restored population are taken to establish multiple new independent populations; and a ‘stepping-stone’ approach, where founders are sequentially introduced from newly recovered populations. This work represents the first analytical comparison of effectiveness between reintroduction strategies and aims to provide general reintroduction recommendations for endangered species that require careful management for population expansion. Our second additional analysis explores parameter space beyond that of the crested ibis, to investigate more generally how demography and inbreeding interact to impact population viability in restoration programs.

## Results

### Model overview

The simulation model sets out from a bottlenecked population consisting of a few individuals that form long-term breeding pairs through random mating (Fig 1A). Each pair reproduces once per year. Their reproductive success is determined by the species’ clutch size, hatching success (which is influenced by the offspring’s inbreeding coefficients), and fledging success of the hatchlings. An individual’s inbreeding coefficient is calculated by tracking a pedigree matrix that contains all individuals that have ever appeared in the population (see Methods for details). After reproduction, individuals survive with probabilities depending on their life stage (i.e., juvenile or adult). If both partners of a breeding pair survive, they will maintain the partnership and breed in the next season. If one of the partners dies or reaches the maximal age of reproduction, the other will randomly form a new pair with an unpaired adult individual of the opposite sex. This generates a population of overlapping generations. Simulations are terminated when the population reaches a threshold size or goes extinct.

**Figure 1.**
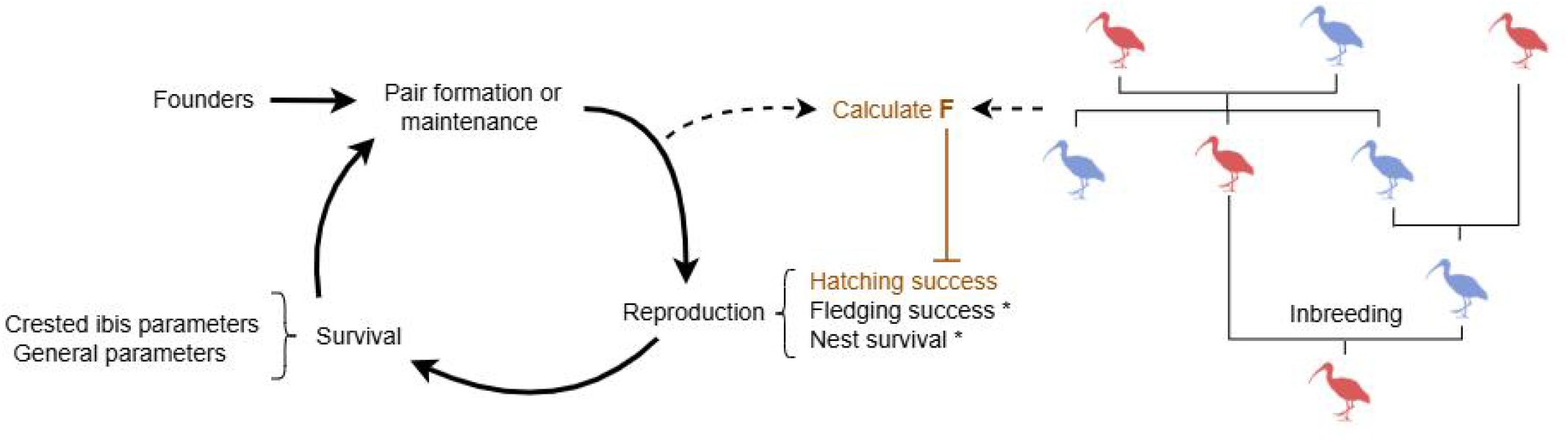
Schematic diagram of the model. In all modeling scenarios, population restoration begins with a small founder population. The breeding cycle consists of pair formation or maintenance, reproduction, and survival stages. The inbreeding coefficients (*F*) of a breeding pair’s offspring can be calculated by tracking the pedigree (exemplified by the simple pedigree on the right, see Methods for details). Birds of different colors in the cartoon pedigree represent individuals of different sexes. In our model, inbreeding was considered to only negatively impact hatching success. Fledging success, nest survival, and other parameters relevant to survival (see Table 1 for details) are included in the crested ibis model and the two reintroduction model variants (marked by asterisks), but not for the general model analysis that explore broader parameter ranges.

### Theoretical predictions align well with the empirical crested ibis population restoration

To investigate the empirical restoration of the crested ibis, we incorporated its species-specific biological information into the model by analyzing the empirical data (See the online repository^33^). The information includes the kinship of the founder population, the negative correlation between hatching success and inbreeding coefficient based on analyzing the empirical pedigree, the clutch size, the fledging success and nest survival, and stage-specific mortality rates (for details see Methods and Table 1). Out of a total of 300 simulations, 292 replicates (97%) resulted in successful population recovery (Fig. 2A, Supplementary Fig. 1), suggesting that the empirical success of ibis restoration is largely repeatable. Furthermore, the average restoration time for successful simulated restorations (mean±sd: 43.53 ± 8.26 years) and the average population inbreeding coefficients in the final restoration year (mean±sd: 0.16 ± 0.02, Supplementary Fig. 2) both align closely with the empirical situation (restoration time: 41 years; grey dashed line in Fig. 2A; average inbreeding coefficients of one sampled population: mean±sd: 0.17±0.05, grey dashed line in Supplementary Fig. 2) in the crested ibis system. This suggests that our model provides a good fit for the real-world dynamics of the restored crested ibis population. Among simulations that led to successful restoration, most populations entered exponential growth after a short initial fluctuation, whereas in the unsuccessful simulations, population sizes remained small, never entered steady exponential growth, and eventually decreased to extinction mostly within a few years (Fig. 2B).

**Table 1.** Parameters used in the individual-based models.

| Parameter | Description | Value |
| --- | --- | --- |
| (A) breeding parameters in the crested ibis model and the reintroduction model |  |  |
| $C$ | Clutch size | 3 |
| $s_0$ | Coefficient of inbreeding depression | 1 |
| $H_0$ | Baseline Hatching success | 0.9 |
| $R_0$ | Fledging success | 0.62 |
| $R_n$ | Nest survival rate | 0.62 |
| $m_{j1}$ | First year juvenile mortality rate | 0.22 |
| $m_{j2}$ | Second year juvenile mortality rate | 0.05 |
| $m_a$ | Adult mortality rate | 0.05 |
| (B) parameter ranges in the general model |  |  |
| $m_{p_a}$ | Adult mortality rate | 0.0 - 0.5 with an interval of 0.05 |
| $m_{p_j}$ | Juvenile mortality rate | 0.0 - 0.9 with an interval of 0.05 |
| $C_p$ | Clutch size | 1 – 6 |

**Figure 2.**
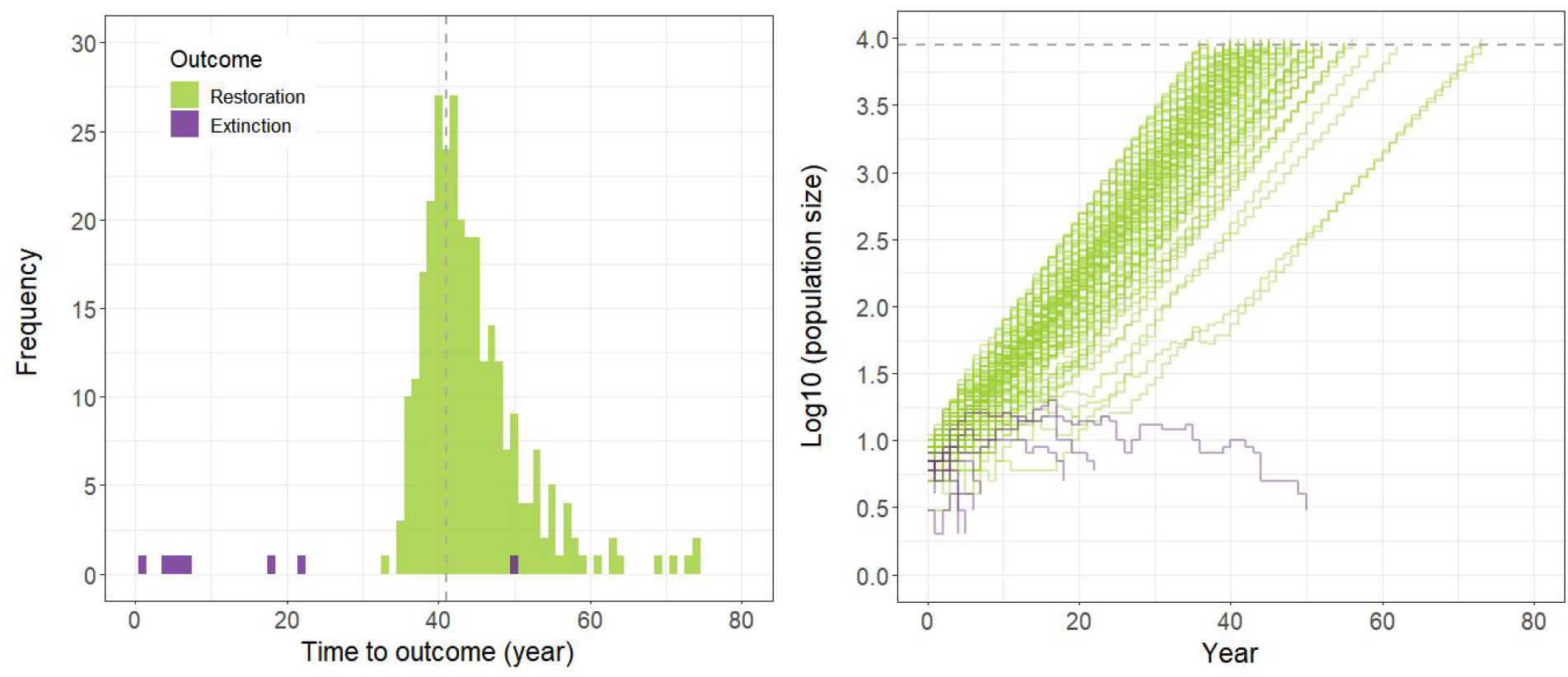
Simulation outcomes of crested Ibis restoration. (A) the distribution of the time a population took to achieve successful restoration (in green, n= 292 replicates) or go extinct (in purple, n=8 replicates) out of 300 simulation replicates. The gray dashed line represents the empirical time (41 years) the crested ibis population took to recover from seven individuals to over 9000. (B) Simulated population dynamics (plotted in the Log_10_ scale) started from the bottlenecked scenario with seven individuals (i.e., two breeding pairs and three juveniles). The 100 trajectories presented here include all eight replicates that ended in extinction and 92 randomly selected replicates that led to successful restoration (see Supplementary Fig. 1 for trajectories of all 300 replicates).

### ‘Firework’ is a more effective reintroduction approach than ‘stepping-stone’

After showing that our theoretical model fits well with the empirical crested ibis restoration, we conducted further model analyses to draw broader conclusions from the successful example. The first analysis includes two model variants, which capture two applied reintroduction strategies in conservation programs, a ‘firework’ and a ‘stepping-stone’ approach (Fig. 3). Continuing with the ibis model, we randomly sampled the same number of individuals (varied from 3 to 13 breeding pairs) from the restored population as founders to simulate the two reintroduction approaches such that their long-term outcomes can be compared. We simulated each reintroduction scenario repeatedly and found that the ‘firework’ approach consistently led to high restoration success across different sizes of founder populations, with only one simulation replicate resulting in extinction (in the scenario with three founding pairs; Supplementary Fig. 3A). We exemplify simulation outcomes in Fig. 3A by showing that the population average inbreeding coefficient initially increases but subsequently reaches a plateau. The average inbreeding coefficient is lower, and the restoration time is shorter for simulations started with more founders (also see Supplementary Fig. 3A). The initial increase in the population-level inbreeding coefficient was due to unavoidable inbreeding caused by the small size of the founder population. Once the population survives this initial phase, individuals that are less inbred can produce more offspring, contributing to an increase in population size and a decline in the variance of inbreeding coefficients.

**Figure 3.**
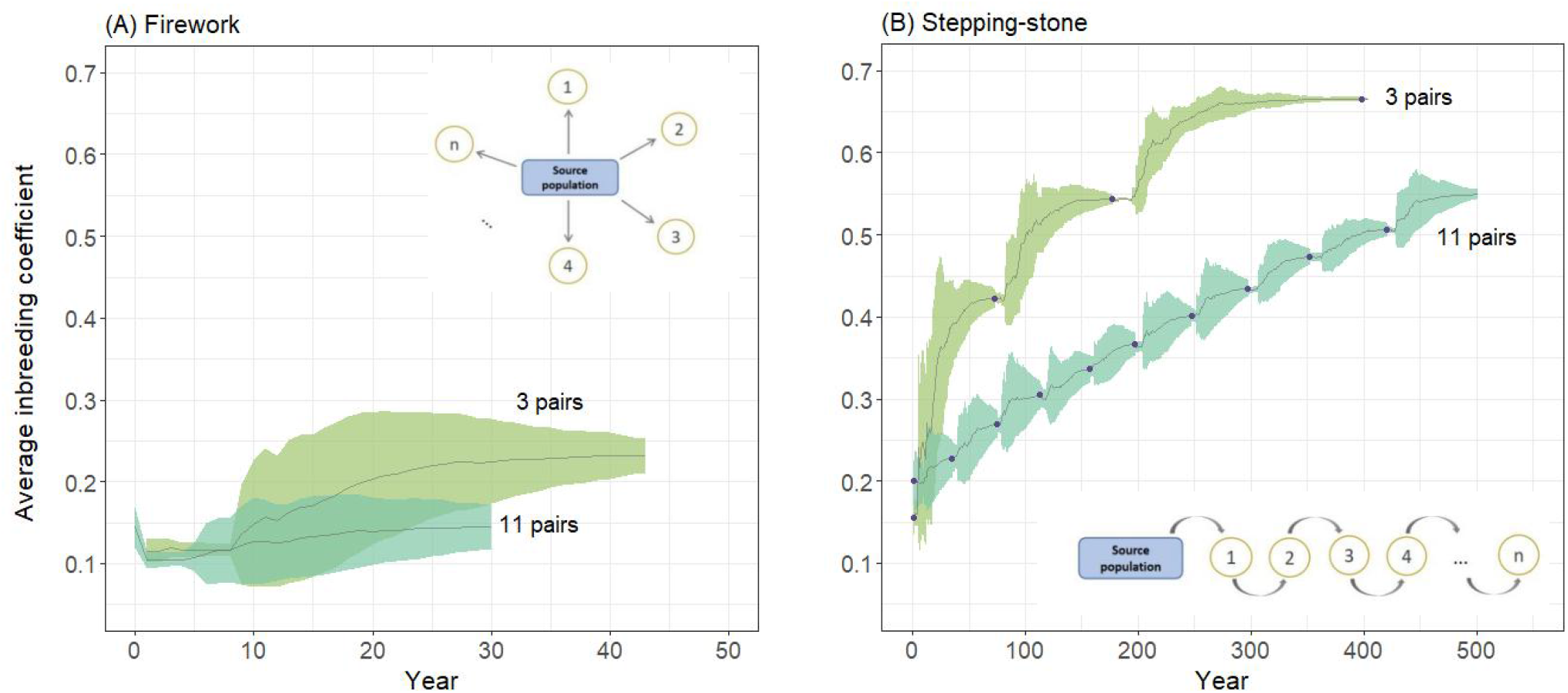
Comparing the effectiveness of the ‘firework’ and ‘stepping-stone’ reintroduction approaches. A conceptual illustration of each approach is presented in the inset of the corresponding panel. One random simulation replicate from different scenarios is used to demonstrate the modeling outcomes. (A) Simulated trajectories where individuals are reintroduced following the ‘firework’ approach with 3 or 11 founder pairs. The gray solid lines represent the trajectory of the population-average inbreeding coefficient (*F*); the colored ribbons depict the standard deviation of *F* over the course of simulations. (B) Simulated trajectories where individuals are reintroduced following the ‘stepping-stone’ approach with 3 or 11 founder pairs. The purple dots indicate the mean *F* of the founders at the beginning of each reintroduction round. The simulation with three founder pairs collapsed at the beginning of the 4^th^ round, while the one with 11 founder pairs achieved all ten rounds of restoration.

We found a similar pattern that the population-level inbreeding coefficient first increases and then stabilizes in each consecutive reintroduction following the ‘stepping-stone’ approach. However, there is a notable accumulation of inbreeding coefficient over subsequent re-introduction rounds (Fig. 3B). This approach, therefore, results in more frequent population reintroduction failures, especially when starting with a small number of founder pairs (Supplementary Fig. 3B). For instance, with only three founder pairs, sequential reintroduction following the ‘stepping-stone’ approach led to dramatically increased inbreeding coefficients at the population level, resulting in a sudden population collapse at the beginning of the fourth reintroduction round (Figure 3B). In contrast, simulations starting from 11 founder pairs all led to ten rounds of successful reintroduction. In general, the full program of ten reintroduction rounds was accomplished only when more than seven pairs were introduced as founders in each round following the ‘stepping-stone’ approach (Supplementary Fig. 3B). Our simulations showed that, for both reintroduction approaches, increasing the number of founders leads to faster population growth and mitigates the accumulation of inbreeding coefficients over reintroduction steps. The more founders reintroduced, the smaller difference on the restoration effectiveness between approaches. But overall, the ‘firework’ approach outperforms the ‘stepping-stone’ approach in terms of higher probability, shorter time, and fewer founder pairs required for successful reintroduction.

### The net effect of inbreeding depression on population restoration depends on demography

We further analyzed the model to explore how inbreeding and demography interact to influence population viability under broader parameter space of population demography. Specifically, we varied the clutch size, juvenile mortality rate, and adult mortality rate within ranges that were chosen based on the demographic parameters of 41 long-lived bird species (Supplementary Table 1). We ran simulations with and without the negative impact of inbreeding on the hatching success. The comparison helped us disentangle the interactions between demographic parameters and inbreeding depression on population viability after a bottleneck event.

From a full analysis across these parameter combinations, we can categorize species into three types (markers I, II, and III in Figures 4A-C). Type I species (of which the crested ibis is an example) have favorable demographic rates (e.g., low juvenile and adult mortality rates). For these species, inbreeding depression has a negligible impact on the restoration success after a bottleneck, as they have a high recovery probability regardless (Fig. 4A-C). In contrast, type III species have unfavorable demographic rates and correspondingly a low recovery probability regardless of the impact of inbreeding depression (Fig. 4A-C). Type II species, however, have demographic rates that result in a high chance of population restoration when inbreeding is assumed to be neutral (Fig. 4A), but have a low probability of recovering from a bottleneck under inbreeding depression (Fig. 4B-C). All the three types are reflected by the distribution of the 41 species in the full simulation analysis (Fig. 5), where variations in clutch size introduce more complex influences on restoration success. Specifically, small clutch sizes considerably hinder species with high adult mortality from recovery, whereas population restoration in species with larger clutches is mostly impeded by high juvenile mortality. Although inbreeding depression is assumed to act uniformly across model scenarios, the net impact of inbreeding on population recovery from a bottleneck varies with demographic rates. Given the large variation in avian demography (Supplementary Table 1), our model results highlight the necessity to consider the interactions between a species’ demography and the impact of inbreeding depression when evaluating the likelihood of population recovery from bottleneck events.

**Figure 4.**
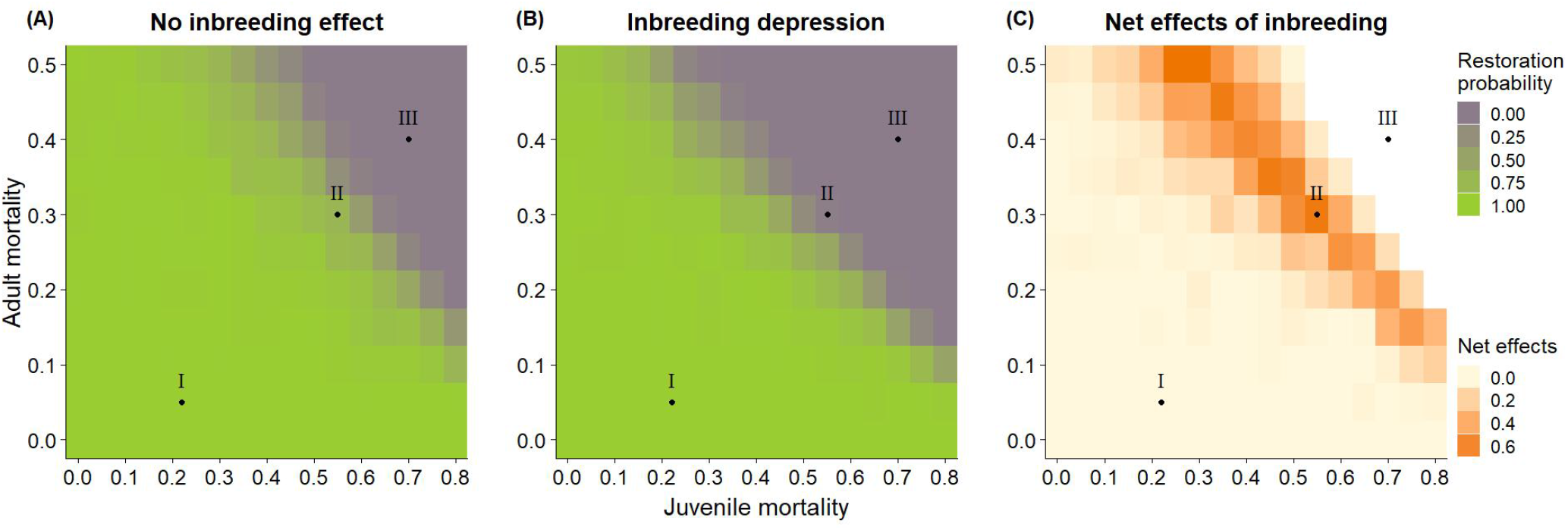
Disentangling the impact of inbreeding depression on population recovery in a broad demographic parameter space. The outcomes were drawn from simulations started with a bottlenecked founder population of three pairs and a scenario where the clutch size was set to three eggs. The value in each cell was calculated based on 100 independent simulation replicates. The heatmaps in (A) and (B) show the probability of population restoration under broad ranges of juvenile and adult mortality rates when the effect of inbreeding depression is respectively *absent* or *present*. The color scale is consistent in both panels. (C) The net effect of inbreeding depression on the probability of population restoration. Each cell in the grid represents the restoration probability difference between the corresponding cells in (A) and present (B). The white area presents a parameter space where restoration is not possible regardless of the presence or absence of inbreeding depression. The labeled dots I,II and III in the panels represent hypothetical species whose restoration probabilities are influenced by the combined effect of inbreeding depression and demographic parameters in different ways.

**Figure 5.**
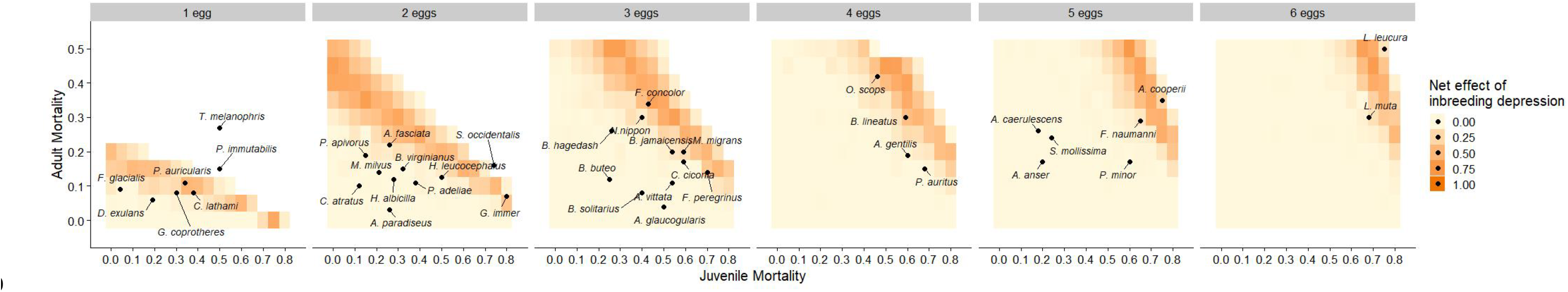
The net effect of inbreeding depression on population recovery across broad parameter ranges of juvenile and adult mortality. 41 avian species (in black dots, see Supplementary Table 1 for the full Latin name) are mapped in the panels according to their demographic rates (i.e. juvenile and adult mortality rates and clutch size) to predict possible effects of inbreeding depression on the potential of restoration of a bottlenecked population under our model assumptions. The colored gradient, ranging from light to dark orange, represents an increasing net impact of inbreeding depression on the probability of population restoration. Population restoration was not possible in the white areas.

## Discussion

We developed an individual-based model inspired by the empirical restoration of the crested ibis, and performed two additional analyses to draw broader conclusions from this remarkable conservation success. Our base model provides a good fit to the ibis example, as demonstrated through the agreement between simulation results and empirical data in terms of population restoration time and average inbreeding coefficient. On this basis, we further adjusted and analyzed the model to compare the outcomes of two applied reintroduction strategies and found the ‘firework’ approach to be more effective than the ‘stepping-stone’ approach. The former approach generally requires fewer founders and leads to a higher restoration probability, shorter restoration time, and lower inbreeding levels. We subsequently analyzed the model to investigate beyond the demographic parameters of the crested ibis and demonstrate that a bottlenecked population’s chance of recovery depends on the interplay between its vulnerability to inbreeding depression and its species-specific demography.

Inbreeding can pose a significant extinction risk to bottlenecked populations, such as during the early stages of reintroduction schemes. Our simulations show that the ‘firework’ approach outperforms the ‘stepping-stone’ approach by mitigating the accumulative effect of inbreeding that occurs through sequential reintroduction rounds. There is empirical evidence for the success of ‘firework’ schemes, as seen in the white-tailed deer (*Odocoileus virginianus*) and bearded vultures (*Gypaetus barbatus*), with reintroduced populations maintaining steady growth over time^7, 31^. However, this approach is not guaranteed to always lead to success, as demonstrated by the conservation project of the Seychelles Magpie-robin (*Copsychus sechellarum*), which was successful in some translocation attempts but failed in others^9^. The varied mortality risks induced by local ecological threats are proposed to play a role, reflecting the importance of demography in determining restoration success. The disadvantage of the ‘stepping-stone’ approach revealed in our study aligned with another theoretical model^34^, which showed a dramatic increase in extinction rate after the second (compared to the first) bottleneck event. Empirically, sequential bottlenecked populations introduced following the ‘stepping-stone’ approach have been found to suffer from reduced genetic diversity, such as several second-order bottlenecked populations of the New Zealand Saddleback (*Philesturnus carunculatus rufusater*) and the dice snakes (*Natrix tessellata*) illegally introduced into the lakes north of the Alps in Switzerland^35, 36, 37^. In practice, reintroduction programs often combine the two approaches. For example, the reintroduction of the Alpine ibex (*Capra ibex*) started with the ‘firework’ approach to establish the first three wild populations in Switzerland^6^. After that, some secondary reintroductions continued to follow the ‘firework’ approach, while others adopted the ‘stepping-stone’ approach with founders taken from a single reintroduced population. Genetic analysis, aligned with our model conclusion, showed a stronger loss of genetic diversity in the secondary reintroduced populations following the latter approach. Our work underscores that theoretical analysis can help clarify ambiguous outcomes in empirical reintroduction programs and make predictions to optimize conservation strategies.

From a practical conservation perspective, our simulation outcomes suggest that inbreeding-induced restoration failure is most likely to occur during the early phase of population establishment. In successful cases, both the population restoration and reintroduction models exhibit similar dynamic patterns of the population-level inbreeding coefficient (*F*). The rapid initial increase in *F* indicates severe inbreeding occurs due to the small population size. The average *F* later reaches a plateau once the population overcomes the first hurdle, accompanied by converging individual variation of *F* over the course of time. This trend indicates that the population enters a positive eco-evolutionary vortex^38^, where inbreeding probability declines with the chance of encountering close kin in an expanding population. These findings provide insights for conservation evaluation, especially for the recovery state of closed populations or captive restoration programs.

Our model also helps to further understand the effects of inbreeding on population viability in general. It is often puzzling that many populations show steady growth despite being marked by known negative effects of inbreeding on reproductive fitness and decreased genetic diversity^16, 25, 26, 27^. Taylor *et al*. (2017)^16^ emphasized that population growth is not a reliable criterion for predicting population viability, as genetic load created by inbreeding in the spotted kiwi (*Apteryx owenii*) has resulted in reduced hatching success, which was predicted to eventually breakdown the population. Similarly, despite rapid population recovery, concerns arose after genetic analyses confirmed high levels of inbreeding in the endangered black-faced spoonbill (*Platalea minor*)^25^. Results from our model suggest that considerable inbreeding does not necessarily hinder populations to recover from a bottleneck, especially for species with high survival and reproductive rates. Yet, populations with high mortality rates will similarly likely face high extinction risks due to this unfavorable demography, with thus little impact of inbreeding depression, as for example seen in the continuous failed restoration of the great bustard (*Otis tarda*), given its generally low survival rates^8^. Meanwhile, the negative effects of high mortality rates and inbreeding depression on population recovery can sometimes be compensated by large clutch sizes, showing the complex roles of different demographic traits in influencing population viability. In general, our results highlight that the impact of inbreeding on population viability is dependent on the species’ demography, and thus both aspects should be carefully considered when predicting population viability or attempting reintroduction schemes.

As demonstrated here, theoretical models offer a powerful and feasible approach for analyses of empirical population restorations and to be able to draw broader conclusions from them, information which itself can be used to help inform future conservation efforts. Whilst our current model is tailored towards a crested ibis population, we encourage future models to use inbreeding depression and demographic data from other species to draw more specific population restoration predictions. For instance, in alignment with empirical evidence from the crested ibis, we here assume inbreeding negatively affects embryo development, but in other species inbreeding depression may affect different aspects of fitness. Other factors can also be incorporated into such models, either to make more species-specific predictions or to gain more general understanding of the range of factors that can influence population viability^39^. For example, inbreeding-induced genetic purging^4, 40^ or Allee effects^41, 42^ could both affect the restoration success of small populations. Allee effects have been found to vary across populations of the crested ibis depending on local geographic features and habitat quality^43^, highlighting that incorporating spatial structure and environmental heterogeneity into future models can also be informative. Despite the simplified assumptions, our simulations across a broad demographic space reveal a crucial but often overlooked role of demography in shaping population viability, which can potentially signal species vulnerability under current model premise. We encourage empirical studies to collect more quantitative data on species-specific inbreeding effects and demographic information, in combination with theoretical analyses such as the one presented here, that together can provide useful insights into the development of successful conservation and population restoration programs.

## Methods

### Calculating inbreeding coefficients from simulated pedigree

The inbreeding coefficient *F* measures the level of inbreeding, which is the probability that two alleles at a given locus in an individual are identical by descent from a common ancestor. This can be estimated using pedigrees to track the kinship of all common ancestors of an individual’s parents. We calculate inbreeding coefficients by generating a numerator relationship matrix (***A***) using the tabular method^43^. Specifically, all individuals in the pedigree are numbered from 1 to *N* and stored in a *N* × *N* matrix containing the pairwise relationship coefficients between any two individuals. The diagonal element corresponding to a focal individual *i* is calculated by 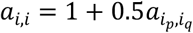. Here, *i*_*p*_ and *i*_*q*_ represent the parents of the focal individual *i*, located in the *p*-th and *q*-th row of the matrix; and 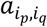 stands for the relationship coefficient between the parents (Supplementary Methods). In the lower triangular matrix, the remaining values in the *i*-th row can be calculated by 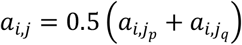. Following this method, all elements of ***A*** can be calculated through pedigree tracking. The inbreeding coefficient of individual *i* is then derived from its corresponding diagonal element in the ***A*** by *F*_*i*_ = *a*_*i, i*_ − 1 . We programmed the pedigree matrix and the calculation of *F* in C++ with an efficient method that majorly reduces the storage space required by the matrix. The pseudocode of the algorithm is provided in the Supplementary Methods and the C++ script is accessible in the Dryad repository^33^.

### Model simulation details for crested ibis restoration

We parameterized the individual-based model based on the breeding system and other life history data of the crested ibis. Simulations were initiated with a small population of seven individuals from two families (two pairs and three nestlings), resembling the population state when the species was rediscovered in 1981 in the wild^28^. Pedigree analysis of one sampled population revealed that inbreeding mainly and negatively influences the brood hatching success, but showed little influence on fertilization success, nestling survival and fecundity (relevant data and analyses are provided in the Dryad repository^33^). The average number of hatchlings *V* is determined by the clutch size *C* and hatching success, which is negatively correlated with the inbreeding coefficients (in Eq. 1, *s* = *s*_0_). The parameter values are provided in Table 1.

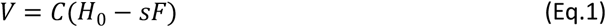

The number of fledglings was drawn from a Poisson distribution with a mean value of *VR*_0_, where *R*_0_ reflects the fledging success of the hatchlings. We also consider the empirical nest survival rate *R*_*n*_, which reflects the probability a nest survives the breeding season without damage or predation. As crested ibises remain in the juvenile stage without breeding for two years, mortality rates were parameterized as three stages based on empirical data: first-year juvenile mortality rate, second-year juvenile mortality rate, and adult mortality rate. The survivorship of juveniles and adults in the population after a breeding season is determined by sampling a Bernoulli distribution with their corresponding life-stage-dependent mortality rates as the mean. Survivors enter a new breeding cycle, except for the first-year juveniles. Simulations were terminated once the population reached 9000 individuals to compare the simulated population recovery with the empirical restoration outcomes.

### Model analysis 1: comparing the ‘firework’ and ‘stepping-stone’ reintroduction approaches

Based on the crested ibis model, we developed a reintroduction analysis that includes two model variants to compare the conservation effectiveness of two applied reintroduction strategies, the ‘firework’ approach and the ‘stepping-stone’ approach. In each independent simulation of the ‘firework’ approach, we first generated a source population containing over 9000 individuals using the previously introduced crested ibis model. From the same source population, a fixed number of founder pairs were randomly sampled 10 times to initialize 10 rounds of reintroduced populations. Each reintroduction round was terminated either when the population recovered to over 5000 individuals or went extinct. The number of founder pairs ranged from 3 to 13 across simulation scenarios, each was simulated 10 times independently (Figure 3A). In each independent simulation of the ‘stepping-stone’ approach, the founders were always taken from the most recently recovered population to form a series of reintroductions (Figure 3B). The number of founder pairs also ranged from 3 to 13. Each reintroduction round proceeded to the next one once the population size reached 5000. Simulations were terminated either at the end of 10 rounds of successful reintroduction or when a population went extinct in one round. We ran 50 independent simulation replicates for each number of founder pairs.

### Model analysis 2: evaluating the impact of inbreeding on general species-related demographic parameters

To evaluate the net effects of inbreeding depression on the probability of population recovery in a bottleneck scenario, we implemented two model variants where the effect of inbreeding on hatching success was switched on (*s* = *s*_0_ = 1) or off (*s* = 0). We also explored broader parameter ranges of the mortality rate of juveniles 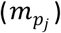 and adults 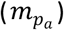 and clutch size (*C*_*p*_) beyond that of the crested ibis (Table 1). We implemented a factorial combination of 11 different 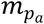 values, 17 different 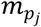 values, and 6 different *C*_*p*_ values. Each unique parameter combination was simulated 100 times independently. This model was initiated from a small population consists of 3 non-related pairs. The linear correlation between inbreeding coefficients and hatching success was kept the same due to the limited availability of relevant data for other species. To draw more general conclusions beyond the crested ibis example, we did not consider nest failure or hatchling death, and all individuals older than one year were allowed to form breeding pairs. Finally, we mapped 41 avian species that we considered long-lived and have empirical data from the COMADRE Database (2024) (Available from: https://www.compadre-db.org/ [15 July 2024, Version 4.23.3.1]) to the simulation outcomes to provide implications of varied inbreeding effects on species with diverse demographic rates under our model assumptions.

## Acknowledgments

We acknowledge funding from the National Natural Science Foundation of China (Young Scientists Fund no. 32301290 to J.Z.); Chinese Academy of Sciences (CAS Pioneer Hundred Talents program) and the Institute of Zoology, Chinese Academy of Sciences (2023 and 2024) to D.W.; and the Swiss National Science Foundation (Grant no. 211549 to X-Y.L.R.). We thank Stefano Canessa for insightful discussions. We also thank the Centre for Information Technology of the University of Groningen for their support and for providing access to the Hábrók high-performance computing cluster.

