## supplement for "Inbreeding and demography interact to impact population recovery from bottlenecks"

This Supplementary document includes:

**Supplementary Methods.** An efficient algorithm for calculating the inbreeding coefficients

**Supplementary Fig. 1.** Complete results of the simulated temporal dynamics of the crested ibis population size.

**Supplementary Fig. 2.** The temporal dynamics of the population-level inbreeding coefficient of the crested ibis model.

**Supplementary Fig. 3.** Comparisons of the restoration probability, time and average *F* between two reintroduction approaches.

**Supplementary Fig. 4.** The net effect of inbreeding depression on population recovery across broad parameter ranges of juvenile and adult mortality.

**Supplementary Table 1.** Demographic information of 41 long-lived bird species.

**Supplementary Methods: An efficient algorithm for calculating the inbreeding coefficients**

Tier (1990)[1] introduced a matrix-based algorithm to calculate inbreeding coefficients.

The inbreeding coefficient $F$ of the $i$-th individual is given by its corresponding diagonal element in the pedigree matrix $A$:

$$F_i = a_{i,i} - 1. \qquad \text{(Eq. S1)}$$

The matrix is populated recursively by tracking the relationship between a focal individual and all others in the population.

$$\begin{cases} a_{i,j} = 0.5\left(a_{i,j_p} + a_{i,j_q}\right) \\ a_{i,i} = 1 + 0.5a_{i_p,i_q} \end{cases}, \qquad \text{(Eq. S2)}$$

where the subscripts $i$ and $j$ refer to individuals $i$ and $j$ (with $i < j$, since the matrix is symmetrical); $j_p$ and $j_q$ refer to the parents of individual $j$. For individuals with at least one unknown parent (assumed to be unrelated, marked with an asterisk),

$$a_{i_{p^*},i_q} = a_{i_p,i_{q^*}} = a_{i_{p^*},i_{q^*}} = 0. \qquad \text{(Eq. S3)}$$

Eq. S3 implies that for all founders, the diagonal element $a_{i,i} = 1$ and the inbreeding coefficient $F_i = 0$, meaning no inbreeding is considered between founders in our models. Figure M1 shows an example of populating the $A$ matrix of a small population of 8 individuals by tracking the corresponding pedigree.

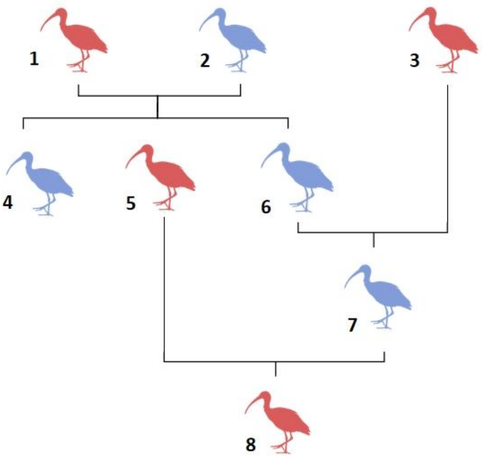


| | 1 | 2 | 3 | 4 | 5 | 6 | 7 | 8 |
|---|---|---|---|---|---|---|---|---|
| **1** | 1 | | | | | | | |
| **2** | 0 | 1 | | | | | | |
| **3** | 0 | 0 | 1 | | | | | |
| **4** | 0.5 | 0.5 | 0 | 1 | | | | |
| **5** | 0.5 | 0.5 | 0 | 0.5 | 1 | | | |
| **6** | 0.5 | 0.5 | 0 | 0.5 | 0.5 | 1 | | |
| **7** | 0.25 | 0.25 | 0.5 | 0.25 | 0.5 | 0.5 | 1 | |
| **8** | 0.375 | 0.375 | 0.25 | 0.375 | 0.75 | 0.5 | 0.75 | 1.125 |

**Supplementary Method Fig. 1.** An example of a simple pedigree and the corresponding matrix used to calculate inbreeding coefficients.

Tier (1990) noted that not all entries in the matrix are required to calculate the required inbreeding coefficients in the population, and that a recursive algorithm can be used to calculate the required coefficients, avoiding unnecessary calculations. Whereas Tier made use of a linked list, here, we make use of a std::map object in C++, which is a lookup table for any $a_{i,j}$, based on $i$ and $j$. This lookup table helps us avoid repeating calculations of $a_{i,j}$ entries. Furthermore, we require three vectors to hold crucial information: a vector of the indices of all mothers ($\boldsymbol{p}$), a vector of the indices of all fathers ($\boldsymbol{q}$), and a vector of all $a_{i,i}$ values (i.e., the diagonal entries in the matrix). We use the following pseudocode to illustrate how to obtain arbitrary $a_{i,j}$ values efficiently.

```
CREATE lookup_table
CREATE mother
CREATE father
CREATE diagonal

FUNCTION calculate_a_i_j (arguments: i, j )

    LET ANSWER 0
    LOOKUP aij IN lookup_table with arguments i, j
    IF FOUND: SET ANSWER = entry IN lookup_table

    IF (i == 0 OR j == 0) SET ANSWER = 0
    IF (i == j) SET ANSWER = diagonal at position i

    LET m = mother AT position j
    LET f = father AT position j


    LET a_ip = calculate_a_i_j (m, i)
    LET a_iq = calculate_a_i_j (f, i)

    SET ANSWER 0.5 + (a_ip + a_iq)

    ADD 1 TO lookup_table
    RETURN ANSWER
```

Upon the birth of a new individual $i$, we need to update the diagonal vector by calculating the corresponding $a_{i,i}$ element to ensure the algorithm works properly, and thus, we call:

```
LET m = mother AT position i
LET f = father AT position i

LET a_pq = calculate_a_i_j(m, f)
SET diagonal at position k = 1 + 0.5 * a_pq
```

The $a_{i,i}$ values of the founders were initialized to take the value 1 at the start of the simulations.

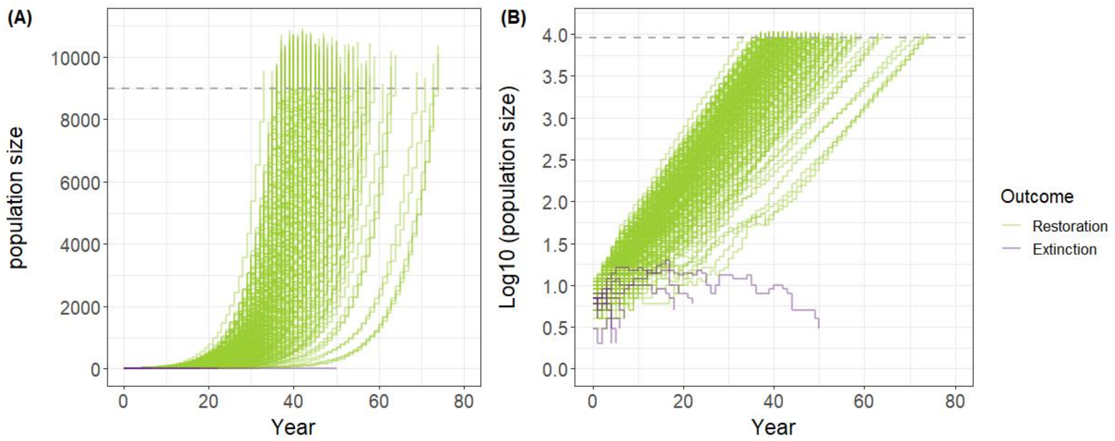


**Supplementary Fig. 1. Complete results of the simulated temporal dynamics of the crested ibis population size.** (A) shows the population size trajectory over time steps in all 300 replicates; (B) presents the log-transformed (with a base of 10) population size trajectory over the course of simulations. Successful population restorations (n = 292) are depicted in green, and failed ones (n = 8) are in purple. The gray dashed lines indicate the threshold population size (absolute or log-transformed 9000 individuals) that terminated the simulation runs.

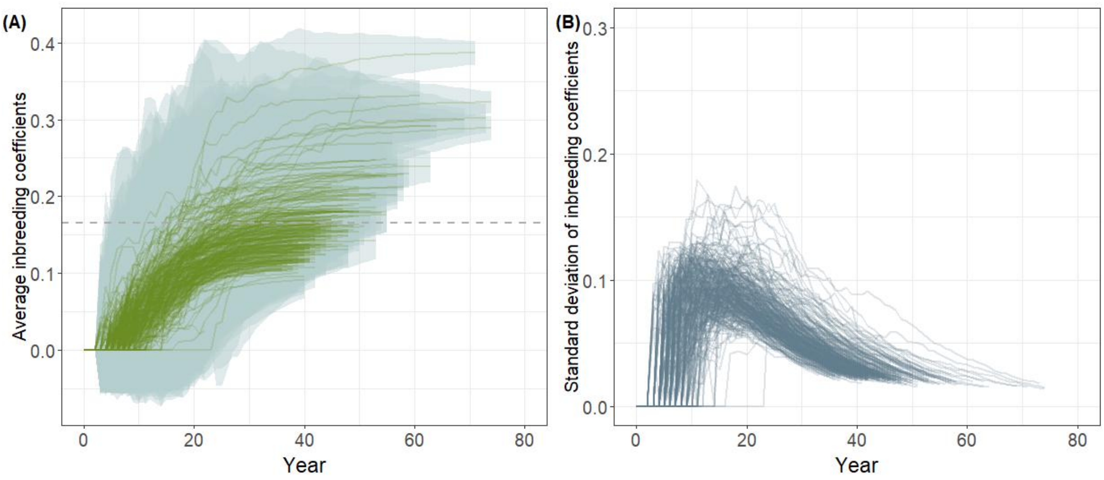


**Supplementary Fig. 2.** The temporal dynamics of the population-level inbreeding coefficient of the crested ibis restoration model. (A) The trajectories of population-level average inbreeding coefficients ($F$) across 300 simulation replicates. Dark green solid lines represent the dynamics of population average $F$, while the width of light blue ribbons depicts the standard deviation of $F$. The gray dashed line indicates the average $F$ of the current (until 2023) crested ibis population from empirical data. (B) Dynamics of the standard deviation of $F$ at the population level over the course of simulations. Each gray trajectory corresponds to one independent simulation replicate.

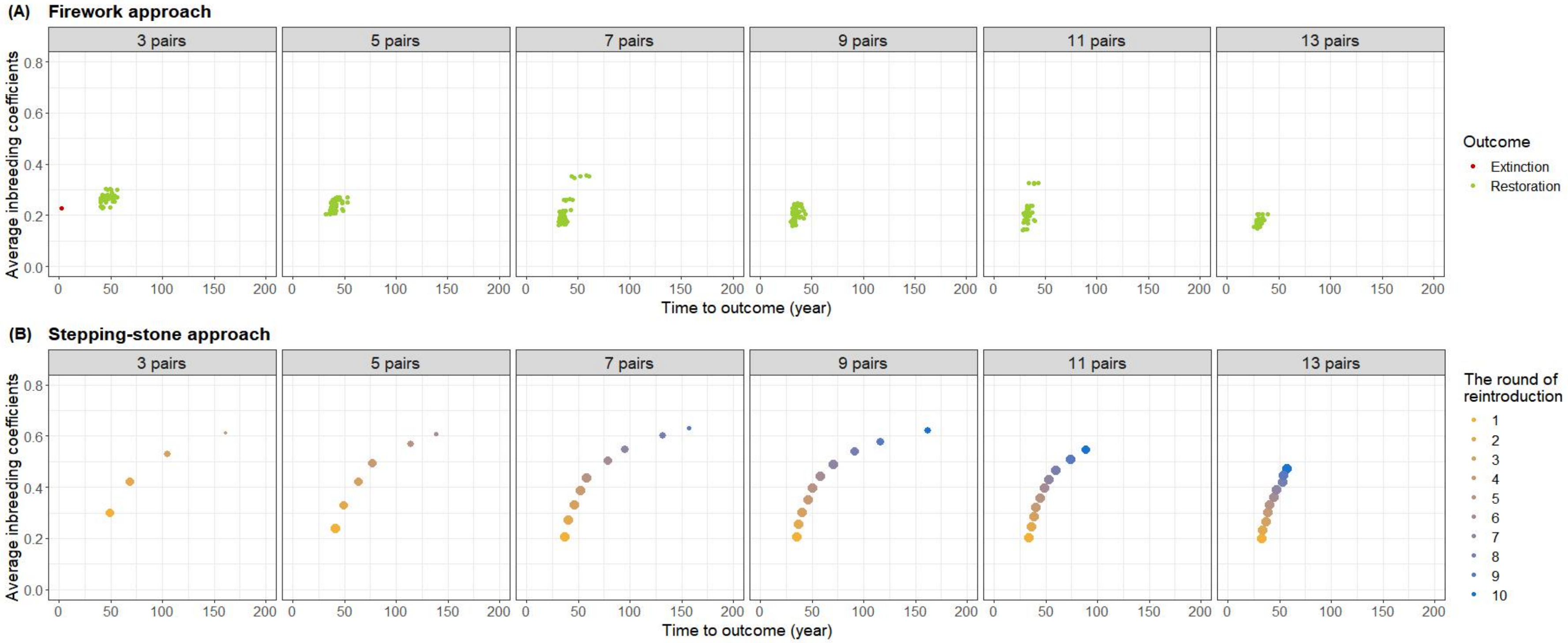


**Supplementary Fig. 3.** Comparisons of the restoration probability, time and population-average $F$ (at the end of the simulations) between the 'firework' and 'stepping-stone' reintroduction approaches. The subpanels in each row represent 6 simulation scenarios where the number of founders ranged from 3 to 13 pairs; each scenario was simulated 50 times independently. (A) displays a series of simulation outcomes using the 'firework' approach. Green dots in each panel represent successful restorations in simulation runs. The red dot in the first subpanel (reintroducing with 3 founder pairs) represents a simulation run that led to population extinction. (B) shows the simulation outcomes for the 'stepping-stone' reintroduction approach. The number of successful consecutive reintroduction rounds is depicted by different colors. The dot size reflects the number of reintroductions that succeeded in specific round out of the 50 simulation runs, with larger dots indicating more times of successful restorations.

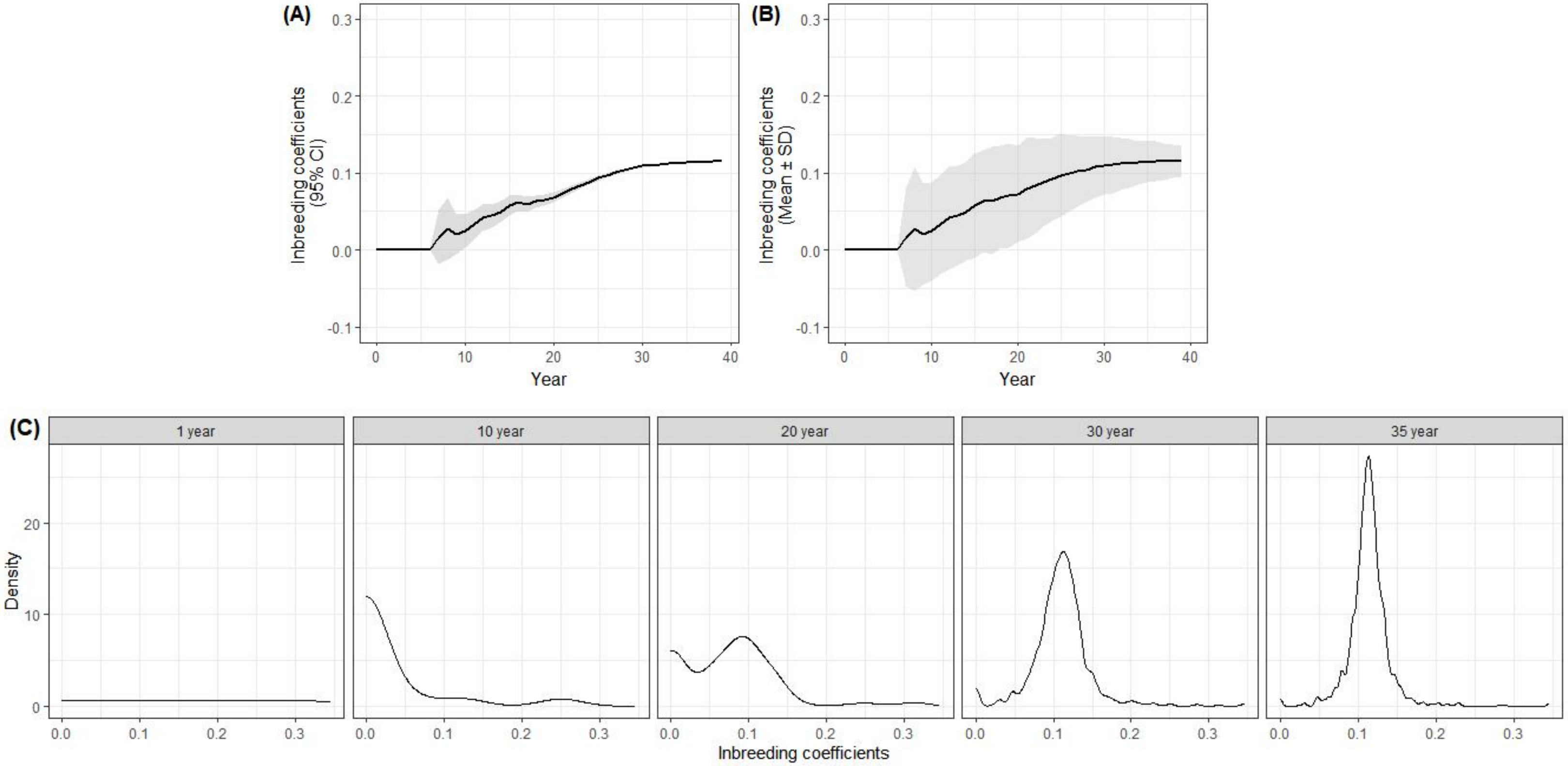


**Supplementary Fig. 4.** (A–B) Comparing two ways of presenting the variation of inbreeding coefficients at the population level for the same population dynamics trajectory. The gray ribbon represents the 95% confidence interval and the standard deviation of the inbreeding coefficients in panels (A) and (B), respectively. (C) Snapshots of the distribution of inbreeding coefficients at several time points during the simulation.

**Supplementary Table.** Demographic information of the 41 long-lived avian species.

| Latin name | Short Latin name (used in Figure 5) | Species common name | Juvenile annual mortality rate | Adult annual mortality rate | Average clutch size | Rounded clutch Size |
|---|---|---|---|---|---|---|
| *Gyps coprotheres* | *G. coprotheres* | Cape vulture | 0.3 | 0.08 | 1 | 1 |
| *Diomedea exulans* | *D. exulans* | Wandering albatross | 0.19 | 0.06 | 1 | 1 |
| *Thalassarche melanophris* | *T. melanophris* | Black-browed albatross | 0.5 | 0.27 | 1 | 1 |
| *Phoebastria immutabilis* | *P. immutabilis* | Laysan albatross | 0.5 | 0.15 | 1 | 1 |
| *Fulmarus glacialis* | *F. glacialis* | Northern fulmar | 0.04 | 0.09 | 1 | 1 |
| *Puffinus auricularis* | *P. auricularis* | Townsend's shearwater | 0.34 | 0.11 | 1 | 1 |
| *Calyptorhynchus lathami* | *C. lathami* | Glossy Black Cockatoo | 0.38 | 0.08 | 1 | 1 |
| *Aquila fasciata* | *A. fasciata* | Bonelli's eagle | 0.26 | 0.22 | 1.5 | 2 |
| *Haliaeetus leucocephalus* | *H. leucocephalus* | Bald eagle | 0.5 | 0.125 | 2 | 2 |
| *Haliaeetus albicilla* | *H. albicilla* | White-tailed eagle | 0.28 | 0.12 | 1.7 | 2 |
| *Milvus milvus* | *M. milvus* | Red kite | 0.21 | 0.14 | 2 | 2 |
| *Pernis apivorus* | *P. apivorus* | European honey buzzard | 0.15 | 0.19 | 2 | 2 |
| *Coragyps atratus* | *C. atratus* | Black vulture | 0.12 | 0.1 | 2 | 2 |
| *Bubo virginianus* | *B. virginianus* | Great horned owl | 0.32 | 0.15 | 1.7 | 2 |
| *Strix occidentalis* | *S. occidentalis* | Northern spotted owl | 0.74 | 0.16 | 2 | 2 |
| *Gavia immer* | *G. immer* | Great northern diver | 0.8 | 0.07 | 1.8 | 2 |
| *Anthropoides paradiseus* | *A. paradiseus* | Blue crane | 0.26 | 0.03 | 2 | 2 |
| *Pygoscelis adeliae* | *P. adeliae* | Adelie penguin | 0.38 | 0.11 | 2 | 2 |
| *Milvus migrans* | *M. migrans* | Black kite | 0.59 | 0.2 | 2.4 | 3 |
| *Buteo buteo* | *B. buteo* | Common buzzard | 0.25 | 0.12 | 2.8 | 3 |
| *Buteo jamaicensis* | *B. jamaicensis* | Red-tailed hawk | 0.54 | 0.2 | 2.4 | 3 |
| *Buteo solitarius* | *B. solitarius* | Hawai'ian hawk | 0.4 | 0.08 | 2.4 | 3 |

| | | | | | | |
|---|---|---|---|---|---|---|
| *Falco concolor* | *F. concolor* | Sooty Falcon | 0.43 | 0.34 | 2.4 | 3 |
| *Falco peregrinus* | *F. peregrinus* | Peregrine falcon | 0.7 | 0.14 | 2.8 | 3 |
| *Ciconia ciconia* | *C. ciconia* | White stork | 0.59 | 0.17 | 2.6 | 3 |
| *Amazona vittata* | *A. vittata* | Puerto Rican parrot | 0.54 | 0.11 | 2.8 | 3 |
| *Ara glaucogularis* | *A. glaucogularis* | Blue-throated macaw | 0.5 | 0.04 | 2.4 | 3 |
| *Nipponia nippon* | *N.nippon* | Crested Ibis | 0.22 | 0.05 | 3 | 3 |
| *Bostrychia hagedash* | *B. hagedash* | Hadeda ibis | 0.26 | 0.26 | 2.4 | 3 |
| *Accipiter gentilis* | *A. gentilis* | Northern goshawk | 0.6 | 0.19 | 3.5 | 4 |
| *Buteo lineatus* | *B. lineatus* | Red-shouldered hawk | 0.59 | 0.3 | 3.2 | 4 |
| *Otus Otus scops* | *O. scops* | Eurasian Scops Owl | 0.46 | 0.42 | 3.5 | 4 |
| *Phalacrocorax auritus* | *P. auritus* | Double-crested cormorant | 0.68 | 0.15 | 3.5 | 4 |
| *Accipiter cooperii* | *A. cooperii* | Cooper's hawk | 0.75 | 0.35 | 4.2 | 5 |
| *Falco naumanni* | *F. naumanni* | Lesser kestrel | 0.65 | 0.29 | 4.2 | 5 |
| *Anser anser* | *A. anser* | Greylag goose | 0.2 | 0.17 | 4.9 | 5 |
| *Anser caerulescens* | *A. caerulescens* | Snow goose | 0.18 | 0.26 | 4.5 | 5 |
| *Somateria mollissima* | *S. mollissima* | Common Eider | 0.24 | 0.24 | 4.9 | 5 |
| *Platalea minor* | *P. minor* | Black-faced Spoonbill | 0.6 | 0.17 | 4.9 | 5 |
| *Lagopus leucura* | *L. leucura* | White-tailed ptarmigan | 0.75 | 0.5 | 5.8 | 6 |
| *Lagopus muta* | *L. muta* | Japanese rock ptarmigan | 0.68 | 0.3 | 5.8 | 6 |
